# Transcriptomic and Multi-scale Network Analyses Reveal Key Drivers of Cardiovascular Disease

**DOI:** 10.1101/2024.09.11.612437

**Authors:** Bat-Ider Tumenbayar, Khanh Pham, John C. Biber, Rhonda Drewes, Yongho Bae

**Affiliations:** Department of Pharmacology and Toxicology, Jacobs School of Medicine and Biomedical Sciences, University at Buffalo, Buffalo, NY 14203, USA; Department of Pathology and Anatomical Sciences, Jacobs School of Medicine and Biomedical Sciences, University at Buffalo, Buffalo, NY 14203, USA; Department of Biomedical Engineering, School of Engineering and Applied Sciences, University at Buffalo, Buffalo, NY 14260, USA

**Author notes:** These authors contributed equally to this work (Bat-Ider Tumenbayar, Khanh Pham, John C. Biber).

## Abstract

Cardiovascular diseases (CVDs) and pathologies are often driven by changes in molecular signaling and communication, as well as in cellular and tissue components, particularly those involving the extracellular matrix (ECM), cytoskeleton, and immune response. The fine-wire vascular injury model is commonly used to study neointimal hyperplasia and vessel stiffening, but it is not typically considered a model for CVDs. In this paper, we hypothesize that vascular injury induces changes in gene expression, molecular communication, and biological processes similar to those observed in CVDs at both the transcriptome and protein levels. To investigate this, we analyzed gene expression in microarray datasets from injured and uninjured femoral arteries in mice two weeks post-injury, identifying 1,467 significantly and differentially expressed genes involved in several CVDs such as including vaso-occlusion, arrhythmia, and atherosclerosis. We further constructed a protein-protein interaction network with seven functionally distinct clusters, with notable enrichment in ECM, metabolic processes, actin-based process, and immune response. Significant molecular communications were observed between the clusters, most prominently among those involved in ECM and cytoskeleton organizations, inflammation, and cell cycle. Machine Learning Disease pathway analysis revealed that vascular injury-induced crosstalk between ECM remodeling and immune response clusters contributed to aortic aneurysm, neovascularization of choroid, and kidney failure. Additionally, we found that interactions between ECM and actin cytoskeletal reorganization clusters were linked to cardiac damage, carotid artery occlusion, and cardiac lesions. Overall, through multi-scale bioinformatic analyses, we demonstrated the robustness of the vascular injury model in eliciting transcriptomic and molecular network changes associated with CVDs, highlighting its potential for use in cardiovascular research.

## I. INTRODUCTION

An estimated 127.9 million Americans, or 48.6% of adults aged 20 and above, have some form of cardiovascular disease (CVD) [1], including hypertension and atherosclerosis-associated diseases such as peripheral vascular disease and coronary artery disease. A common etiology of cardiovascular pathologies is the progression of neointimal hyperplasia into atherosclerosis [2], which coincides with arterial stiffening [3] and can lead to cardiac ischemia/infarction, brain ischemia, and thrombosis [4]. Procedures like embolectomy [5], vein grafting [6], balloon angioplasty, and stenting [7] can damage the vessel wall, causing neointimal hyperplasia, restenosis, or thrombosis. Fine-wire vascular injury models are commonly used [8-11] to study the molecular mechanisms of neointimal hyperplasia [12-15]. Neointimal hyperplasia arises from the migration, proliferation, and extracellular matrix (ECM) deposition of vascular smooth muscle cells (VSMC) from the media into the intimal layer, leading to vascular wall thickening and further exacerbating atheroprogression and CVDs. Vascular injury creates conditions that mimic various aspects of CVD, including aberrant proliferation [16], migration [17], differentiation [18-20], ECM synthesis [19], inflammation [21], and loss of cellular contraction [22]. A frequently overlooked feature of the vascular injury model is increased vessel stiffening [23], a mechanosignal that may accelerate neointimal hyperplasia [24-26]. Despite fostering various pathologies associated with CVD in general, vascular injury is not typically used as a model for CVD outside of those that exhibit neointimal hyperplasia and vascular stiffening. Expanding the use of vascular injury model into studying CVD could uncover valuable insights into potential therapeutic targets for treating this comorbidity.

Recent studies reveal a complex interaction between inflammation and the immune response in CVD, suggesting that targeting this response could reduce atherosclerotic events [27, 28]. However, suppressing immune activity increases the risk of infections and other diseases. At the site of vascular injury, macrophages regulate angiogenesis at the vessel wall but also contribute to atherosclerosis by maladaptively promoting further plaque buildup through the accumulation of cells, lipids, and ECM components, thereby worsening CVD [29, 30]. Changes in ECM stiffness and remodeling, in response to vascular injury, have been shown to regulate the tissue repair functionality of macrophages [31], indicating an intricate relationship between ECM modulation and the immune system in CVD. Dissecting this interaction in the context of vascular injury can reveal meaningful molecular targets, interactions, and mechanisms to be further studied as new methods to manage CVD and its pathologies.

While considerable knowledge exists on how the actin cytoskeleton regulates key components of neointimal hyperplasia, including VSMC dedifferentiation [32, 33] and migration [34, 35], the specific changes in the actin cytoskeleton associated with vascular injury remain poorly characterized. Mechanical forces can influence the actin cytoskeleton via well-established integrin-dependent mechanisms that transmit ECM stiffness into actin cytoskeletal arrangements through focal adhesion complexes [36, 37]. Although ECM regulation post-vascular injury is well-understood [25, 38-40], the interplay between ECM and the actin cytoskeleton and its contribution to CVD remains elusive.

Bioinformatic analyses provide insights into the complex interplay often presented in diseases. Once transcriptomic data is obtained, the goal is to understand how biological processes modulate genes and vice versa. Analyses as such reveal how these genes are interrelated, allowing us to establish a hierarchy of pathways that govern the broader biological processes. Multi-scale network analysis can be performed [41, 42] using transcriptomic data [43] to interpret how changes in gene regulation relate to protein-protein interactions (PPI) [44] and their impact on disease progression [45, 46]. This approach also identifies associated biological processes and diseases regulated by differentially expressed genes in a model system. While multi-scale networks are diverse in nature, they generally integrate data to infer biological information across different scales [42, 43, 45, 46]. Transcriptomics provides differential gene expression data from a disease, which can be leveraged by the PPI scale to illuminate protein interactions (communication and networks), as well as post-translational modification and degradation relationships. These insights can then be related to pathways that initiate and drive disease progression.

In this study, we performed multi-scale bioinformatic network analysis using microarray datasets from injured femoral arteries and uninjured contralateral (control) femoral arteries in mice two weeks post-injury to investigate how robust transcriptomic changes in response to vascular injury could potentially affects CVDs. Through Ingenuity Pathway Analysis (IPA) of differentially expressed genes (DEGs) found in our dataset, we identified significant activation of various CVDs such as atherosclerosis, arrhythmia, and vaso-occlusion. Protein-protein interaction (PPI) network formed from DEGs was used to identify seven clusters with distinct functions including, ECM organization, metabolic and biosynthetic processes, immune-related processes, actin organization, and cell proliferation, where most clusters exhibited dense communications with each other. A closer analysis of the communication between the ECM remodeling and immune system or actin reorganization clusters further inferred the effects of vascular injury on modulating the activation of aortic aneurysm, cardiac lesions, cardiac damage, and other diseases.

### II. Transcriptomic and multi-scale network analyses

### A. Differential Gene Expression Analysis

To identify changes in gene expression in healthy and injured mouse arteries, we performed differential gene expression analysis on previously published microarray datasets using the R DESeq2 package [47]. Gene expression changes were calculated as follows:

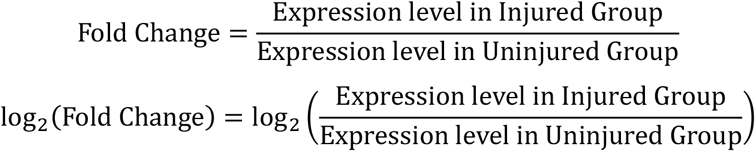

The significance of the results was calculated using the Wald test [47] for p-value calculation and false discovery rate:

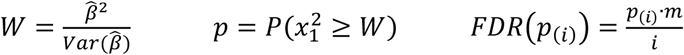

where 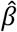 is the estimated coefficient from the regression model, 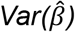 is the variance of the estimated coefficient, 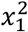 is a chi-square distribution with 1 degree of freedom, *m* is the total number of tests.

### B. Identification of Differentially Expressed Genes

To identify differentially expressed genes (DEGs) in response to vascular injury, the following filtering criteria were applied. Genes (g) were classified as DEGs if they satisfied both of the following conditions:

(i) FDR-adjusted p-value (q-value) threshold: q ≤ 0.15

(ii) log_2_(Fold Change) threshold: ∣log_2_(Fold Change)∣≥ 0.5

Combining these conditions, genes (*g*) are considered significantly differentially expressed if:

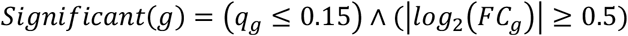

The gene distribution was visualized using a volcano plot created with the Bioinfokit package in Python. The R programming language’s ggplot2 package [48] was used to visualize the Principal Component Analysis (PCA) plot, and covariance was calculated as follows:

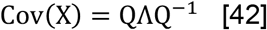

where *Q* is the matrix of eigenvectors and *Λ* is the diagonal matrix of eigenvalues.

### C. Gene Ontology Enrichment Analysis

To explore the biological processes associated with upregulated and downregulated DEGs, gene enrichment analysis was conducted using the g:GOSt function on the gProfiler web server (https://biit.cs.ut.ee/gprofiler/gost) [49]. Given a list of genes G and subsets of upregulated *DEGs G*_*up*.*DEGs*_ and downregulated DEGs *G*_*down*.*DEGs*_ identified by the criteria:

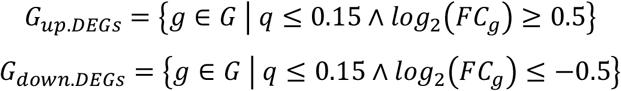

The gene enrichment analysis was then performed using *G*_*up*.*DEGs*_ and *G*_*down*.*DEGs*_ to test for overrepresentation in various gene sets S:

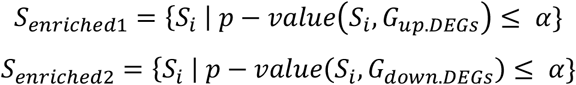

where *S* is the set of all gene ontology (GO) terms being tested, *S*_*i*_ is a particular GO term, *p-value(S*_*i*_, *G*_*DEGs*_) is the statistical significance of the enrichment of *S*_*i*_ in *G*_*DEGs*_, *α* is the significance threshold (*α = 0*.*05*). For visualization purposes, bubble plots representing the top 20 enriched GO terms and KEGG pathways were generated using the SRplot online server.

### D. QIAGEN Ingenuity Pathway Analysis

Combined differential expression analysis results from both clusters 1 and 5, and clusters 1 and 3, were uploaded to the QIAGEN Ingenuity Pathway (IPA) software, using the expression log ratio and p-adjusted values. IPA’s Core Analysis function was employed to investigate altered signaling pathways in response to vascular injury. The Diseases & Functions and Pathways features were used to identify significantly affected pathways and diseases (absolute activation z score ≥ 2; - log(Benjamin-Hochberg p-value ≥ 2) as follows:

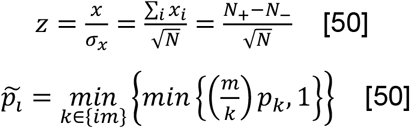

Combining these conditions, a term (*t*) is considered significantly activated or inhibited if:

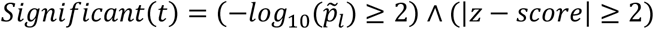

Additionally, Network Analysis feature was used to explore molecular interactions within the combined clusters and their associated diseases and functions. Statistical values for the Network Analysis were computed based on the p-score, derived from p-values and equal to -log10(p-value). The “My pathway” tool was used to illustrate known relationships between molecules or molecules to functions.

To study how molecular-level interactions lead to disease progression, IPA Machine Learning Disease Pathways tool was used to identify similar regulatory patterns among the genes and causally connected them with human diseases. The disease-to-molecule ratio (r) used in IPA Machine Learning Pathways tool was calculated as follows:

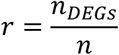

where *n*_*DEGs*_ is the number of DEGs from our dataset that was identified in the pathway, and *n* as the total number of genes that IPA identified in that pathway.

### E. Protein-protein interaction (PPI) network

The STRING website was used to construct the PPI network, and the results were visualized using the Cytoscape software [51]. The expression data for the DEGs were imported into the node table to indicate expression levels using log_2_(fold-change) values and node color to indicate intensity. Orphan and non-present intermediate protein entries were filtered out from the network. K-means clustering tool on the STRING website was used to identify 7 functionally distinct clusters within the PPI network, enrichment analysis for each cluster was conducted using the gProfiler web server.

## III. RESULTS

### A. Genome-wide analysis identifies transcriptomic changes related to CVD in mouse femoral arteries post vascular injury

To investigate the effects of vascular injury on transcriptional responses and biological processes, we performed bioinformatic analyses (**Fig. 1**) on previously published microarray datasets collected from injured and uninjured mouse femoral arteries [52, 53]. Expression values of 21,734 transcripts were identified, and the distinctions among samples (uninjured vs. injured) were visualized in an unsupervised Principal Component Analysis (PCA) plot (**Fig. 2A**). The analysis revealed two distinct clusters of samples, with and without vascular injury, suggesting vascular injury may significantly influence the transcriptomic landscape. To identify differentially expressed genes (DEGs) in our dataset, genes were filtered for q-values of ≤ 0.15 and absolute log2(fold-change) ≥ 0.5. We identified 1,467 DEGs, with 696 upregulated and 771 downregulated. The distribution of these DEGs was displayed in the volcano plot (**Fig. 2B**). To further explore the impact of vascular injury on the biological processes associated with DEGs, we performed Gene Ontology (GO) enrichment analysis. The top 20 biological processes categories enriched among the downregulated DEGs were mainly related to various metabolic/energy and development processes, including “generation of precursor metabolites and energy”, “energy derivation by oxidation of organic compounds”, “system development,” “developmental process,” and “muscle structure development” (**Fig. 2C**). Moreover, the top 20 biological process categories enriched among the upregulated DEGs were primarily related to various biological regulation and cell migration processes, including “positive regulation of biological process”, “response to stress”, and “cell migration” (**Fig. 2D**).

**Figure 1.**
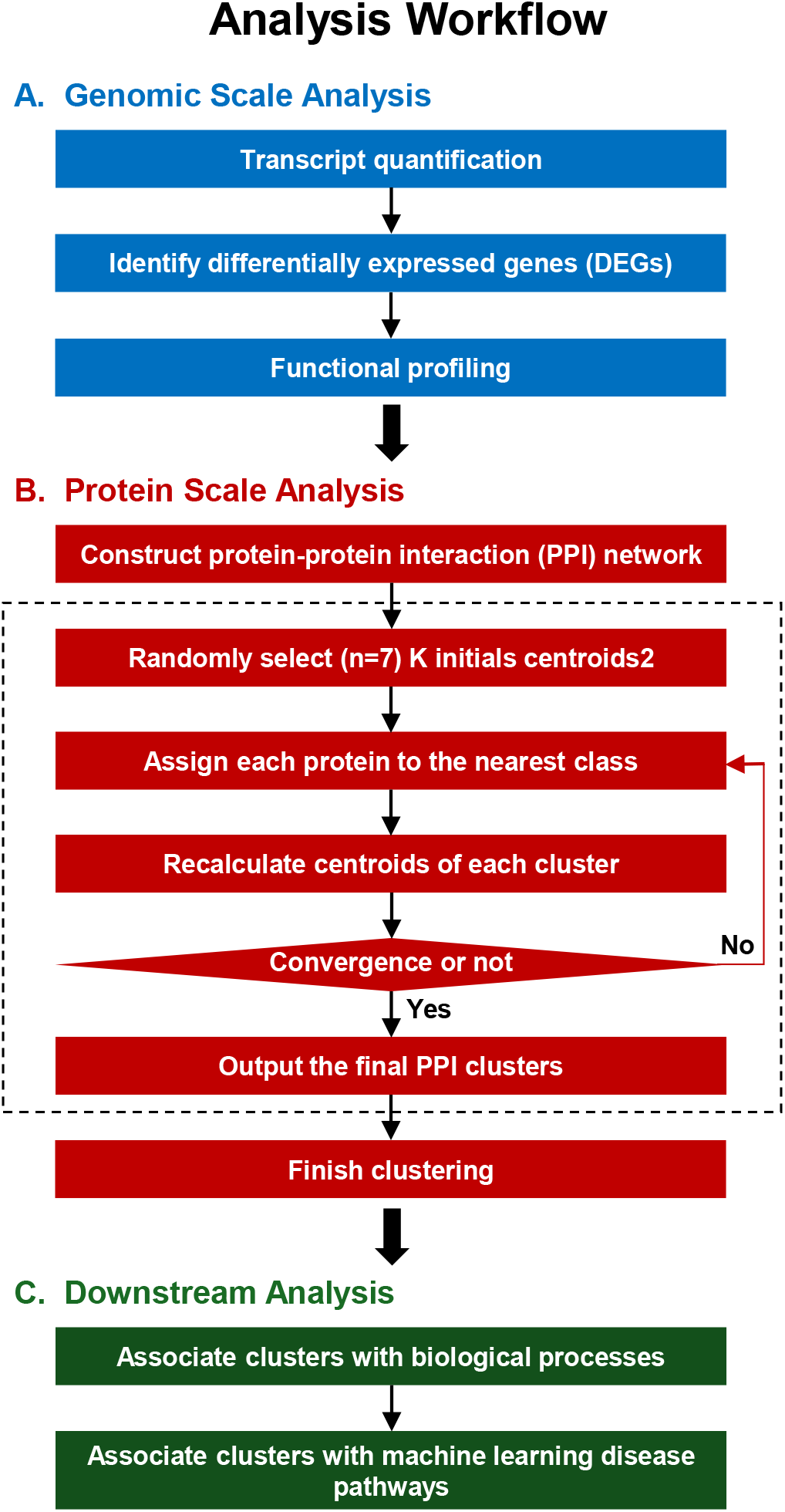
Overview of the multi-scale bioinformatics analysis workflow.

**Figure 2.**
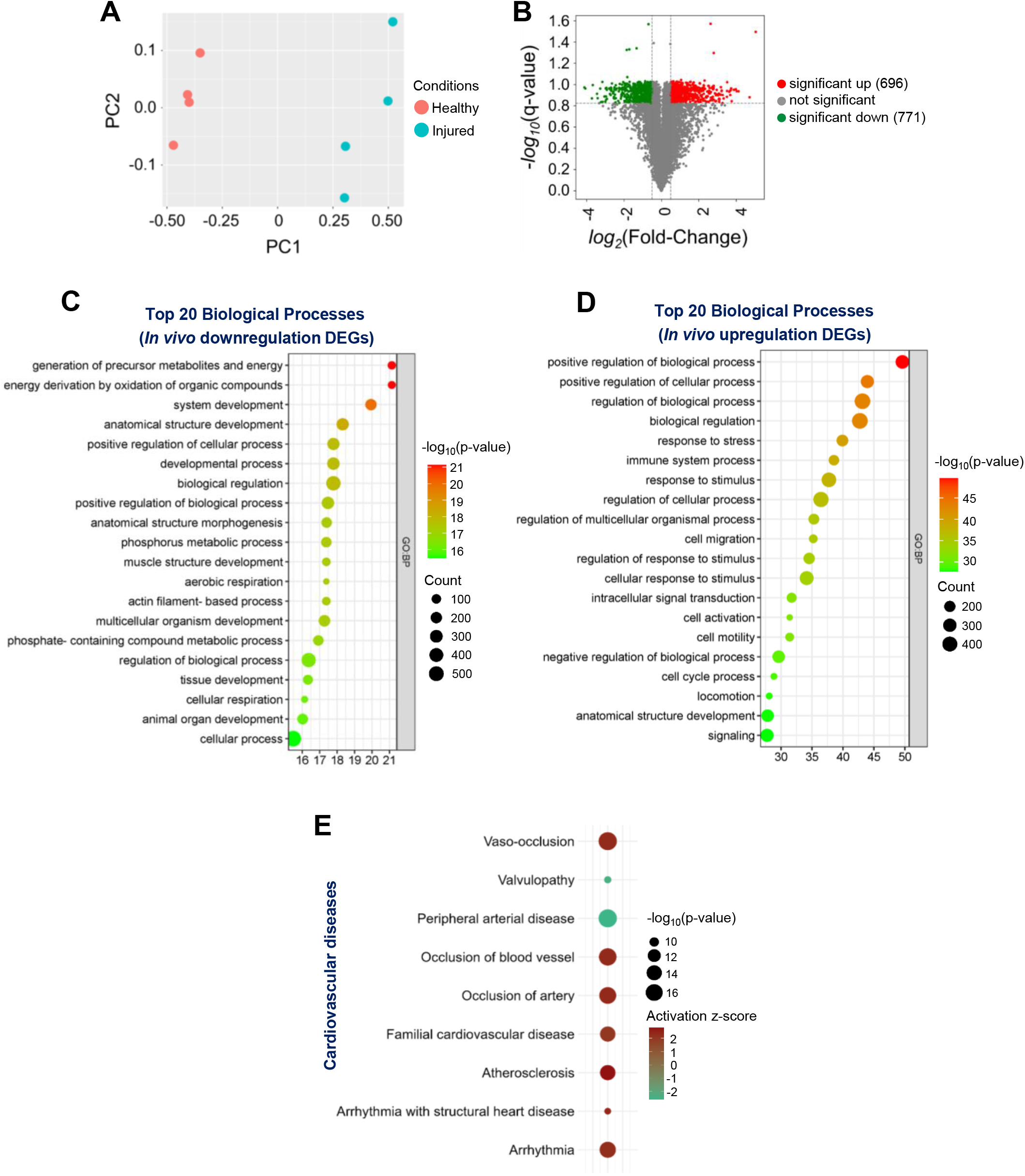
Structure and function of genome-wide transcriptomic changes due to vascular injury. (**A**) Principal Component Analysis (PCA) plot for the entire transcriptome list displays the correlations and variances among the samples. (**B**) Volcano plot illustrates the distribution of differentially expressed genes (DEGs) in response to femoral artery fine-wire injury. Green dots represent statistically downregulated genes (771 downregulated DEGs identified) and red dots represent statistically upregulated genes (696 upregulated DEGs identified). Bubble plots depict the top 20 enriched biological processes for significantly (**C**) downregulated and (**D**) upregulated DEGs. (**E**) Cardiovascular Disease terms were predicted by IPA to be activated in response to vascular injury.

To gain insight into cardiovascular diseases transcriptomically associated with vascular injury, we employed the Core Analysis function of QIAGEN Ingenuity Pathway Analysis (IPA) software on the complete dataset of DEGs (both upregulated and downregulated). Using the IPA Diseases & Functions feature, particularly in the “Cardiovascular Disease” category, seven diseases and functions terms were found to be significantly activated (a Z-score of ≥ 2 is considered significant activation[50], including “Vaso-occlusion” (activation z-score = 2.332), “Arrhythmia” (activation z-score = 2.261), “Atherosclerosis” (activation z-score = 2.772). Two diseases and functions terms were significantly inhibited (a Z-score of ≤ 2 is considered significant inhibition [50] “Peripheral arterial disease” (activation z-score = -2.608) and “Valvulopathy” (activation z-score = -2.401) (**Fig. 2E**). Collectively, these findings indicate that vascular injury markedly alters transcriptomic profiles, thereby modulates a diverse array of cellular behaviors and biological processes, all of which could further the development of CVDs.

### B. Multi-scale analyses identify molecular and functional networks

To integrate the topology information of identified DEGs, a protein-protein interaction (PPI) network was constructed using STRING online database and visualized with Cytoscape software, resulting in 1,188 nodes and 11,025 edges. Further, seven functionally distinct clusters within the PPI network were identified using the STRING online k-means clustering tool (**Fig. 3A**). Cluster 1, consisting of 193 nodes and 533 edges (**Fig. 3B**), was associated with extracellular matrix and development-associated biological processes, including “extracellular matrix organization,” “extracellular structure organization,” “external encapsulating structure organization,” “system development,” “tube development,” and “animal organ development” (**Fig. 3C**). Cluster 2, comprising 177 nodes and 900 edges (**Fig. 3D**), was primarily associated with various metabolic and biosynthetic processes, including “cellular respiration,” “generation of precursor metabolites and energy,” “nucleotide metabolic process,” “purine ribonucleoside triphosphate biosynthetic process,” and “ATP biosynthetic process” (**Fig. 3E**). Cluster 3, consisting of 151 nodes and 1,728 edges (**Fig. 3F)**, was mostly enriched in immune and inflammation-related biological processes, including “immune system process,” “leukocyte activation,” “regulation of immune system process,” “immune response,” and “lymphocyte activation” (**Fig. 3G**). Cluster 4, with 189 nodes and 3,713 edges (**Fig. 3H**), was primarily associated with cell growth, including “cell cycle,” “cell cycle process,” “mitotic cell cycle,” “cell division,” “nuclear division,” and “chromosome organization” (**Fig. 3I**). Cluster 5, consisting of 216 nodes and 584 edges (**Fig. 3J**), was mostly associated with actin cytoskeleton and muscle contraction-related biological processes, including “actin filament-based process,” “muscle system process,” “muscle contraction,” “actin filament-based movement,” “actin cytoskeleton organization,” “cardiac muscle contraction,” and “heart contraction” (**Fig. 3K**). Cluster 6, consisting of 89 nodes and 88 edges (**Fig. 3L**), was mostly associated with various biological regulation processes, including “biological regulation,” “regulation of multicellular organismal process,” “regulation of biological process,” “regulation of hydrolase activity,” “regulation of cell adhesion,” and “regulation of catalytic activity” (**Fig. 3M**). Cluster 7, comprising 45 nodes and 33 edges (**Fig. 3N**), was enriched in various biological processes, including “cellular response to stress,” “DNA damage response,” “regulation of viral processes,” “nucleoside metabolic process,” and “viral process” (**Fig. 3O**). The topological cluster analysis provided significant insights into the distinct biological roles and processes enriched within the protein interactome network, highlighting the extensive transcriptomic changes induced by vascular injury.

**Figure 3.**
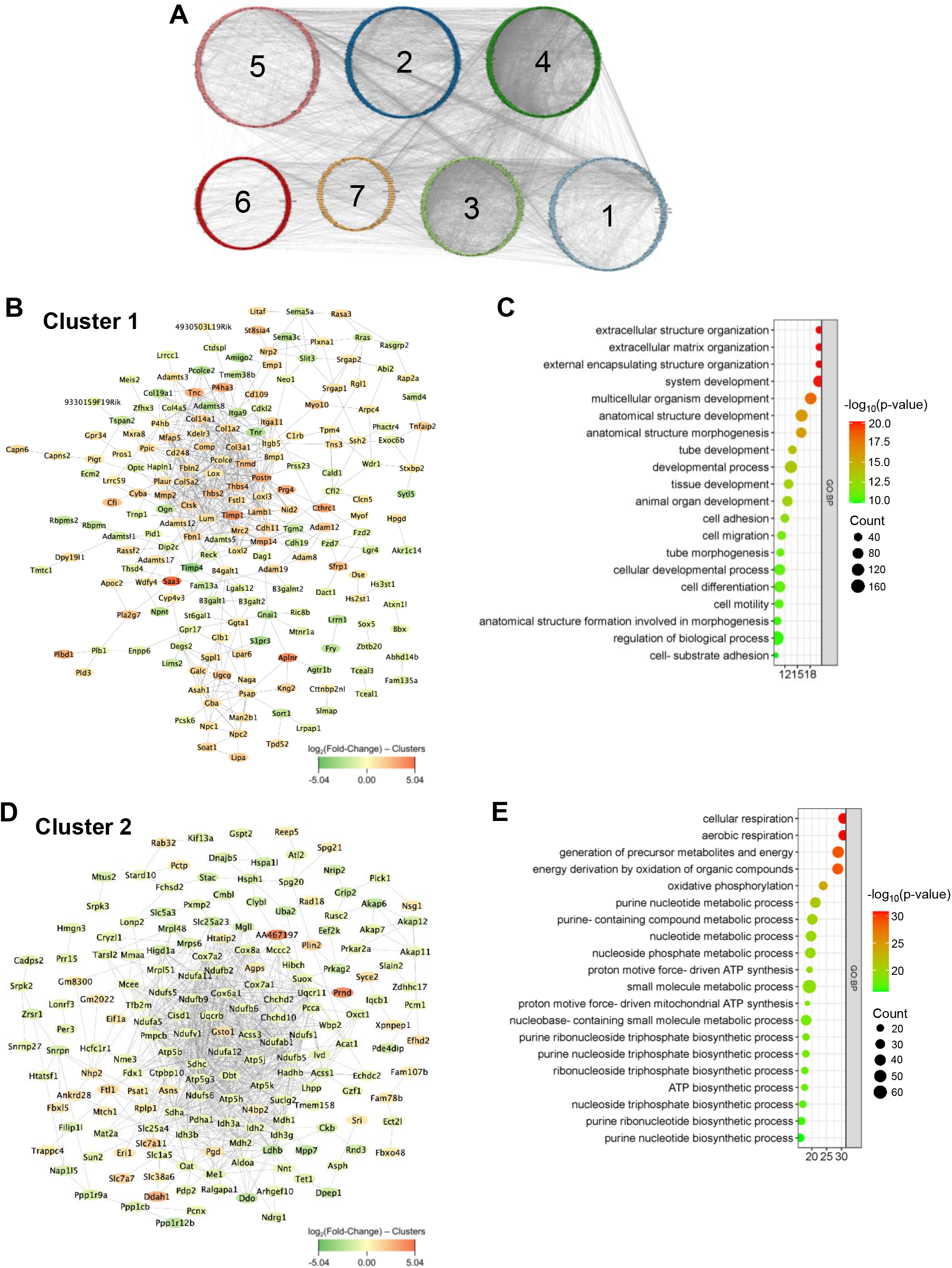
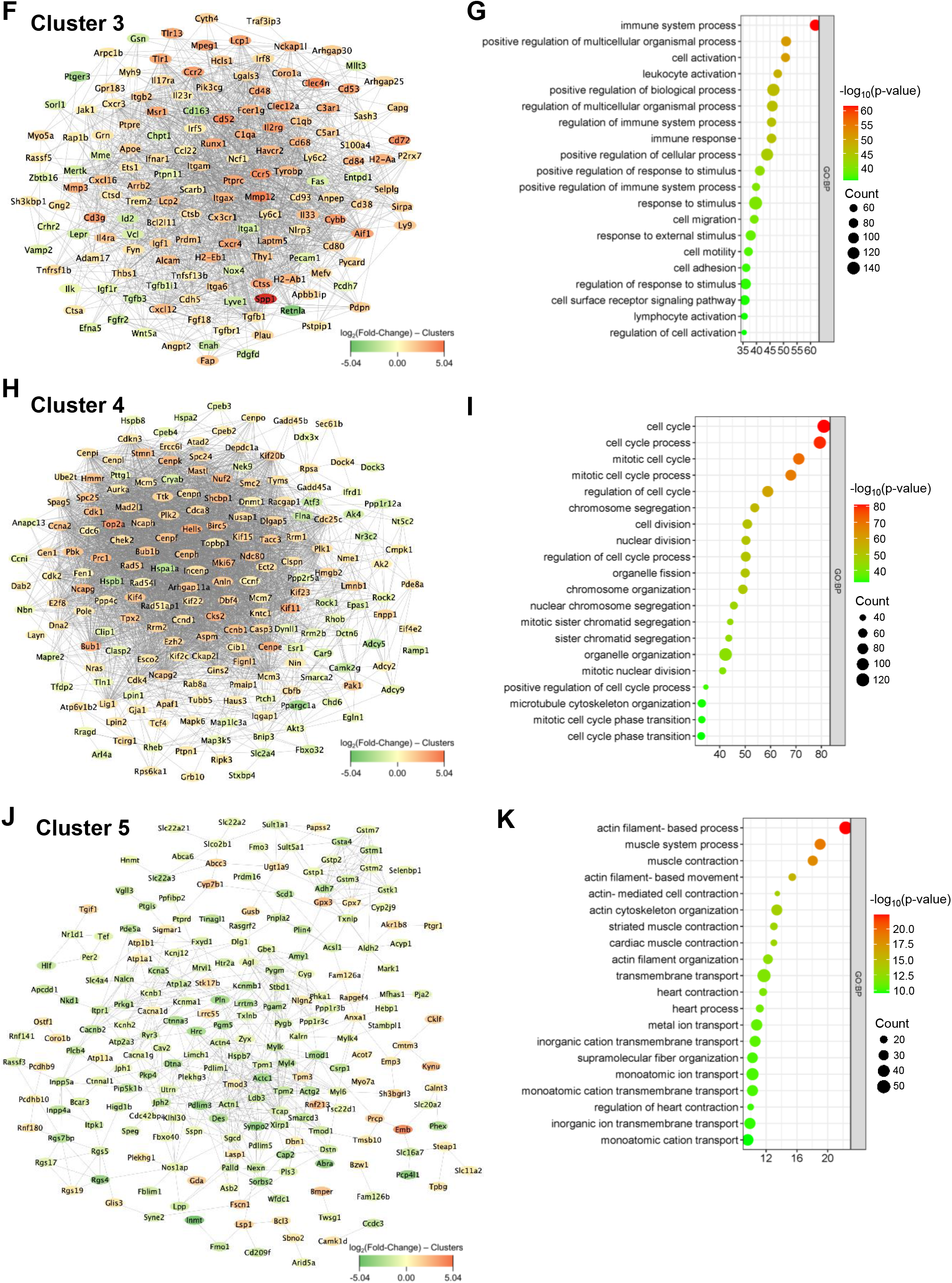
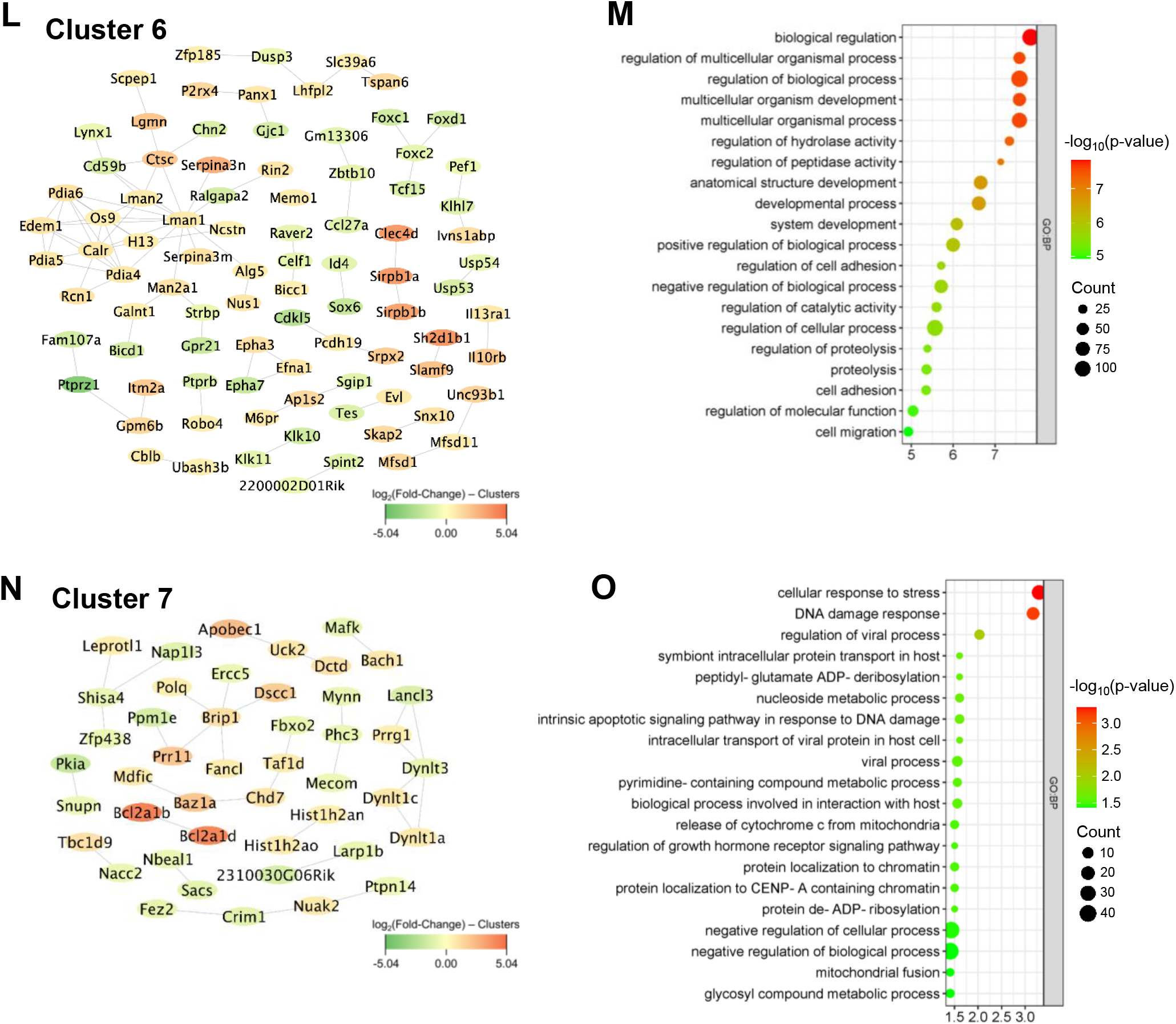
K-means clustering analysis and GO enrichment. (**A**) Network displays 7 clusters within the protein-protein interaction network (1,188 nodes and 11,025 edges) of DEGs based on k-means clustering. (**B-C**) Interaction network of cluster 1 (193 nodes and 533 edges) and its associated biological processes including extracellular matrix organization, extracellular structure organization, and external encapsulating structure organization. (**D-E**) Interaction network of cluster 2 (177 nodes and 900 edges) and its associated biological processes including cellular respiration, aerobic respiration, and generation of precursor metabolites and energy. (**F-G**) Interaction network of cluster 3 (151 nodes and 1,728 edges) and its associated biological processes including immune system process, positive regulation of multicellular organismal process, and cell activation. (**H-I**) Interaction network of cluster 4 (189 nodes and 3,713 edges) and its associated biological processes including cell cycle, cell cycle process, and mitotic cell cycle. (**J-K**) Interaction network of cluster 5 (216 nodes and 584 edges) and its associated biological processes including actin filament-based process, muscle system process, and muscle contraction. (**L-M**) Interaction network of cluster 6 (89 nodes and 88 edges) and its associated biological processes including biological regulation, regulation of multicellular organismal process, and regulation of biological process. (**N-O**) Interaction network of cluster 7 (45 nodes and 33 edges) and its associated biological processes including cellular response to stress, DNA damage response, and regulation of viral process.

### C. Altered molecular communication due to vascular injury contributes to the development of cardiovascular and other diseases

Abnormal remodeling of the actin cytoskeleton and ECM, as well as immune and metabolic dysregulation, and cell overgrowth, ultimately promotes the development of CVDs [30, 54, 55]. Therefore, we assessed the interplay between functionally distinct clusters (**Fig. 3A**) and their combined impact on disease progression by comparing each pair of clusters. Interestingly, the data demonstrated that cluster 3, characterized by an enrichment of immune-related biological processes, exhibited the most significant molecular communications with cluster 1, enriched in ECM structure and organization, and cluster 4, enriched in cell growth (**Fig. 4A**). Additionally, cluster 4 exhibited distinct molecular communications with cluster 2, enriched in metabolic and biosynthetic processes (**Fig. 4A**). Cluster 1 also showed molecular communications with cluster 5, enriched in actin cytoskeleton and muscle contraction-related biological processes (**Fig. 4A**).

**Figure 4.**
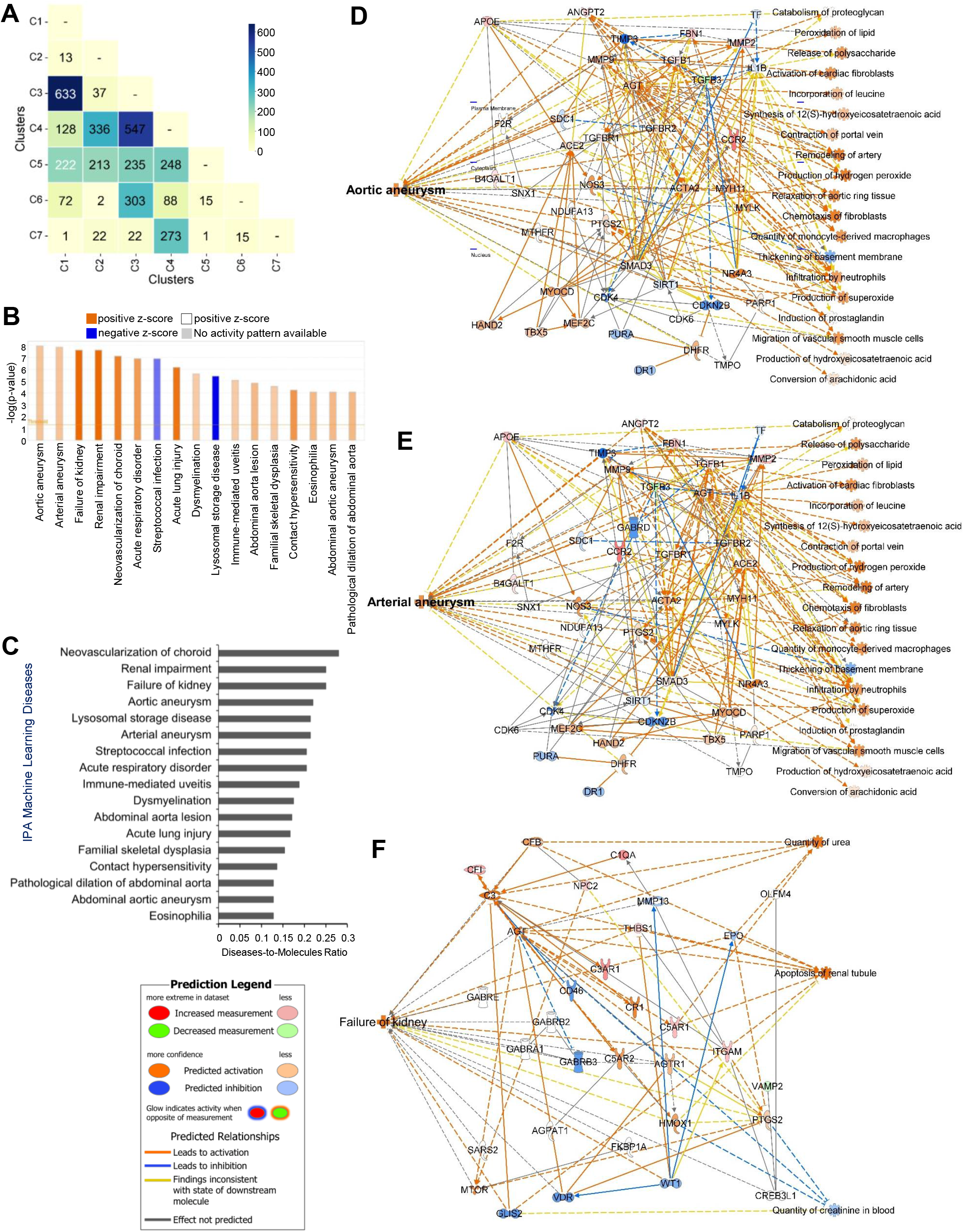
Inter-cluster analysis of diseases and pathways associated with cluster 1 and 3. (**A**) Matrix correlation heatmap illustrates molecular communications between each pair of functionally distinct clusters. (**B**) IPA Core Analysis on cluster 1 and 3 DEGs showing differential changes in various Machine Learning (ML) Disease Pathways. (**C**) Bar plot shows the disease-to-molecules ratio of differentially and significantly changed ML Disease Pathways. Interaction networks display molecular communications and functionalities leading to changes in ML Disease Pathways of (**D**) aortic aneurysm, (**E**) arterial aneurysm, and (**F**) failure of kidney.

We next examined the consequences of molecular communications between cluster 1 (“ECM structure and organization”) and cluster 3 (“immune-related processes”), which had the most significant interactions, using the Core Analysis function and the Machine Learning (ML) Disease Pathways in IPA with the combined DEGs from these clusters. Our findings indicated that pathological vascular conditions, which eventually promote cardiovascular and other diseases, such as “Aortic aneurysm,” “Arterial aneurysm,” “Neovascularization of choroid,” “Abdominal aorta lesion,” “Abdominal aortic aneurysm,” and “Pathological dilation of abdominal aorta” were predicted to be significantly activated (z-score > 4) (**Fig. 4B**). The data also showed significant activation of other diseases such as “Kidney failure,” “Renal impairment,” “Acute respiratory disorder,” “Acute lung injury,” and “Immune-mediated uveitis” (**Fig. 4B**). Additionally, results from the ML Diseases Pathways function similarly showed that “Neovascularization of the choroid” had the highest the disease-to-molecule ratio at 0.28 while “Aortic aneurysm,” “Arterial aneurysm,” and “Failure of kidney” also exhibited higher ratios of 0.22, 0.214, and 0.25, respectively (**Fig. 4C**), inferring the involvement of DEGs from cluster 1 and cluster 3 as key participants in disease development. Furthermore, the ML Disease pathways identified key molecular players and their interaction networks for the three most significant diseases shown in Figures 4B and 4C: aortic aneurysm, arterial aneurysm, and failure of kidney (**Figs. 4D***−***F**). For example, in the aortic and arterial aneurysm pathways shown in Figures 4D and 4E, ACTA2 and MYH11 genes [56-58], whose mutations are known to be associated with these conditions, were significantly connected with other DEGs within the networks and predicted to be activated in response to vascular injury, linking them to aortic and arterial aneurysms. Similarly, in the failure of kidney pathway shown in Figure 4F, AGT and PTGS2 genes, whose mutations are associated with kidney failure [59-61], were predominantly connected with other DEGs and predicted to be activated in response to vascular injury. Interestingly, AGT and PTGS2 genes were also involved in the disease pathways for aortic and arterial aneurysms (**Figs. 4D, E**). The ML Disease generated networks also predicted the activation states of disease-specific etiology. For instance, in Figure 4E, activation of AGT gene is predicted to not only trigger arterial aneurysm, but also activate “Activation of cardiac fibroblasts,” “Remodeling of artery,” and “Infiltration by neutrophils”. Similarly, in Figure 4F, AGT gene activation is predicted to drive “Apoptosis of renal tubule”, a key factor in kidney failure. Taken together, our analysis demonstrates that abnormal remodeling of the ECM, along with immune and metabolic dysregulation, promotes the development of cardiovascular and other diseases by elucidating significant molecular communications between functionally distinct clusters and identifying key molecular players and pathways associated with these conditions.

### D. Changes in ECM constituents and actin cytoskeleton leads to the progression of cardiovascular diseases

We further investigated the implications of molecular communication between cluster 1 (“ECM structure and organization”) and cluster 5 (“actin cytoskeleton”), using the same methods as shown in Figure 4. Of particular interest, our findings unveiled significant and differential activations of several cardiovascular diseases, including “Cardiac damage,” “Occlusion of the carotid artery,” “Cardiac lesions,” and “Congestive heart failure” (**Fig. 5A**). These activations can arise from pathological changes in ECM structure and organization and actin cytoskeleton induced by vascular injury. Additionally, results from the ML Diseases Pathways function showed higher disease-to-molecule ratios of 0.192 for Cardiac damage, 0.138 for Occlusion of carotid artery, and 0.098 for Cardiac lesion (**Fig. 5B**). Additionally, the ML Disease pathways identified critical molecular players and their communication networks for three significant cardiovascular diseases shown in Figures 5A and 5B: cardiac damage (**Fig. 5C**), occlusion of the carotid artery (**Fig. 5D**), and cardiac lesions (**Fig. 5E**). For example, in the cardiac damage and lesion pathways shown in Figures 5C and 5E, DMD, SGCA, SGCB, and SGCG genes [62, 63] associated with these conditions, were significantly connected with other DEGs and predicted to be inhibited in response to vascular injury, linking them to cardiac impairment. Interestingly, in response to vascular injury, PTK2, COL1A2, and FN1 genes, known to be associated with cardiac fibrosis, were densely connected with other DEGs, and their predicted activation link them to cardiac lesion. Additionally, in the occlusion of carotid artery pathway shown in Figure 5D, S100A8, ITGB2, and PTGS2 genes, associated with carotid artery disease [64-67], were predicted to be activated in response to vascular injury. Overall, these robust integrated analyses demonstrate that vascular injury-induced extracellular matrix and actin cytoskeletal alterations profoundly impact diverse cardiovascular diseases.

**Figure 5.**
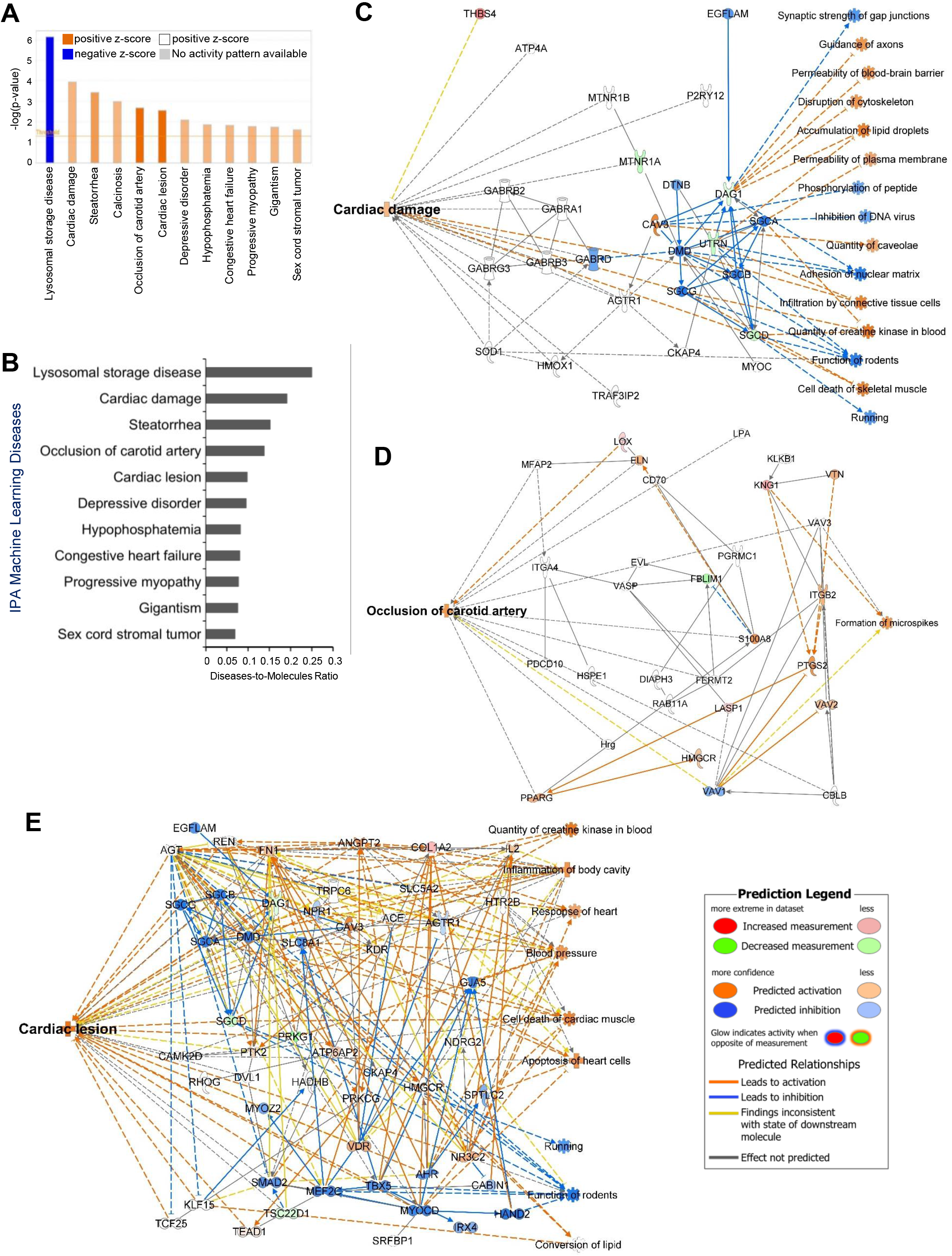
Diseases and pathways associated with cluster 1 and 5. (**A**) IPA Core Analysis on cluster 1 and 5 DEGs showing differential changes in various ML Disease Pathways. (**B**) Bar plot shows the disease-to-molecules ratio of differentially and significantly changed ML Disease Pathways. Interaction networks display molecular communications and functionalities leading to changes in ML Disease Pathways of (**C**) cardiac damage, (**D**) occlusion of carotid artery, and (**E**) cardiac lesion.

## IV. DISCUSSION

In this work, we focused on the biological and molecular scale communications underlying CVD progressions in response to vascular injury. By utilizing bioinformatic sequencing analyses and IPA disease machine learning approaches, we identified complex interactions between DEGs that lead to alterations in biological components, including the actin cytoskeleton, immune system, and ECM. Furthermore, our analysis predicts that interactions among these biological processes and components collectively contribute to the development of various cardiovascular pathologies. Based on the transcriptomic changes revealed by our multi-scale bioinformatic analyses, we suggest expanding the use of vascular injury model as a suitable option to investigate not only neointimal hyperplasia and vessel stiffening, but also a range of other CVDs.

From our DEG list, the IPA Disease and Function feature identified seven CVDs significantly activated in response to vascular injury, including but not limited to vaso-occlusion, atherosclerosis, and arrhythmia. To explore the translational changes due to vascular injury, we constructed a PPI network based on the DEG list and identified functionally distinct clusters within the network. Although distinct, the seven PPI clusters displayed great communications with each other, most significantly between cluster 1 (ECM structure and organization) and cluster 3 (immune-related processes). IPA Disease ML Pathway analysis predicted that crosstalk between these clusters could lead to diseases such as aortic aneurysm, arterial aneurysm, and kidney failure. Our ML analysis also revealed disease-specific networks with key molecular players and etiology. Notably, activation of AGT and PTGS2 gene, known to be associated with kidney failure [59-61], also appeared to influence the aortic and arterial aneurysm networks (**Fig. 4D-F**). Furthermore, interactions between ECM changes and actin cytoskeletal reorganization were linked to cardiac damage, carotid artery occlusion, cardiac lesions, and congestive heart failure. These findings underscore the pivotal roles of ECM and actin cytoskeleton organization alternations in driving vascular pathologies, highlighting the potential relevance of these cellular processes for therapeutic strategies.

## V. CONCLUSION

In conclusion, our study offers a multi-scale level understanding of the intricate regulatory mechanisms governing cardiovascular disease progressions in the context of vascular injury. From genomic level to protein and biological levels, we offered novel insights into the transcriptomic rewiring and molecular networks in response to mouse vascular injury. These findings pave the way for further investigations into the development of targeted therapeutic interventions aimed at modulating ECM, immune response, cytoskeletal dynamics, ultimately contributing to the management and prevention of cardiovascular pathologies.

## ACKNOWLEDGEMENTS

This work was supported by NIH/NHLBI grant No. R01HL163168 to YB.

## AUTHOR DECLARATIONS

The authors have no conflicts to disclose.

## REFERENCES

1. Martin, S.S., et al., 2024 Heart Disease and Stroke Statistics: A Report of US and Global Data From the American Heart Association. Circulation, 2024. 149(8): p. e347–e913.

2. Tang, H.Y., et al., Vascular Smooth Muscle Cells Phenotypic Switching in Cardiovascular Diseases. Cells, 2022. 11(24).

3. Murakami, T., Atherosclerosis and arteriosclerosis. Hypertens Res, 2023. 46(7): p. 1810–1811.

4. Libby, P., et al., Atherosclerosis. Nat Rev Dis Primers, 2019. 5(1): p. 56.

5. Goldberg, E.M., et al., The effects of embolectomy-thrombectomy catheters on vascular architecture. J Cardiovasc Surg (Torino), 1983. 24(1): p. 74–80.

6. Parang, P. and R. Arora, Coronary vein graft disease: pathogenesis and prevention. Can J Cardiol, 2009. 25(2): p. e57–62.

7. Byrne, R.A., et al., Coronary balloon angioplasty, stents, and scaffolds. Lancet, 2017. 390(10096): p. 781–792.

8. Nomura-Kitabayashi, A. and J.C. Kovacic, Mouse Model of Wire Injury-Induced Vascular Remodeling. Methods Mol Biol, 2018. 1816: p. 253–268.

9. Le, V., et al., Murine model of femoral artery wire injury with implantation of a perivascular drug delivery patch. J Vis Exp, 2015(96): p. e52403.

10. Curaj, A., et al., Induction of Accelerated Atherosclerosis in Mice: The “Wire-Injury” Model. J Vis Exp, 2020(162).

11. Lindner, V., J. Fingerle, and M.A. Reidy, Mouse model of arterial injury. Circ Res, 1993. 73(5): p. 792–6.

12. Su, C., et al., Vascular injury activates the ELK1/SND1/SRF pathway to promote vascular smooth muscle cell proliferative phenotype and neointimal hyperplasia. Cell Mol Life Sci, 2024. 81(1): p. 59.

13. Liu, J., et al., C1q/TNF-related protein 4 mediates proliferation and migration of vascular smooth muscle cells during vascular remodelling. Clin Transl Med, 2023. 13(5): p. e1261.

14. Warwick, T., et al., Acute injury to the mouse carotid artery provokes a distinct healing response. Front Physiol, 2023. 14: p. 1125864.

15. Chen, X., et al., Endothelial Foxp1 Regulates Neointimal Hyperplasia Via Matrix Metalloproteinase-9/Cyclin Dependent Kinase Inhibitor 1B Signal Pathway. J Am Heart Assoc, 2022. 11(15): p. e026378.

16. Boehm, M. and E.G. Nabel, The cell cycle and cardiovascular diseases. Prog Cell Cycle Res, 2003. 5: p. 19–30.

17. Zhang, Y., H. Kishi, and S. Kobayashi, Direct active Fyn-paxillin interaction regulates vascular smooth muscle cell migration. J Smooth Muscle Res, 2023. 59: p. 58–66.

18. Li, Y., K.O. Lui, and B. Zhou, Reassessing endothelial-to-mesenchymal transition in cardiovascular diseases. Nat Rev Cardiol, 2018. 15(8): p. 445–456.

19. Prabhu, S.D. and N.G. Frangogiannis, The Biological Basis for Cardiac Repair After Myocardial Infarction: From Inflammation to Fibrosis. Circ Res, 2016. 119(1): p. 91–112.

20. Chen, P.Y., et al., Smooth Muscle Cell Reprogramming in Aortic Aneurysms. Cell Stem Cell, 2020. 26(4): p. 542–557 e11.

21. Elyasi, A., et al., The role of interferon-gamma in cardiovascular disease: an update. Inflamm Res, 2020. 69(10): p. 975–988.

22. Frismantiene, A., et al., Smooth muscle cell-driven vascular diseases and molecular mechanisms of VSMC plasticity. Cell Signal, 2018. 52: p. 48–64.

23. Wang, J., et al., Matrix stiffness exacerbates the proinflammatory responses of vascular smooth muscle cell through the DDR1-DNMT1 mechanotransduction axis. Bioactive Materials, 2022. 17: p. 406–424.

24. Kothapalli, D., et al., Cardiovascular Protection by ApoE and ApoE-HDL Linked to Suppression of ECM Gene Expression and Arterial Stiffening. Cell Reports, 2012. 2(5): p. 1259–1271.

25. Liu, S.L., et al., Matrix metalloproteinase-12 is an essential mediator of acute and chronic arterial stiffening. Scientific Reports, 2015. 5.

26. Mui, K.L., et al., N-Cadherin Induction by ECM Stiffness and FAK Overrides the Spreading Requirement for Proliferation of Vascular Smooth Muscle Cells. Cell Reports, 2015. 10(9): p. 1477–1486.

27. Zhao, T.X. and Z. Mallat, Targeting the Immune System in Atherosclerosis: JACC State-of-the-Art Review. J Am Coll Cardiol, 2019. 73(13): p. 1691–1706.

28. Ridker, P.M., et al., Antiinflammatory Therapy with Canakinumab for Atherosclerotic Disease. N Engl J Med, 2017. 377(12): p. 1119–1131.

29. Chinetti-Gbaguidi, G., S. Colin, and B. Staels, Macrophage subsets in atherosclerosis. Nat Rev Cardiol, 2015. 12(1): p. 10–7.

30. Moore, K.J. and I. Tabas, Macrophages in the pathogenesis of atherosclerosis. Cell, 2011. 145(3): p. 341–55.

31. Meizlish, M.L., et al., Mechanosensing regulates tissue repair program in macrophages. Sci Adv, 2024. 10(11): p. eadk6906.

32. Shi, J.H., J.K. Wen, and M. Han, [The role of SM22 alpha in cytoskeleton organization and vascular remodeling]. Sheng Li Ke Xue Jin Zhan, 2006. 37(3): p. 211–5.

33. Nagayama, K., A Loss of Nuclear-Cytoskeletal Interactions in Vascular Smooth Muscle Cell Differentiation Induced by a Micro-Grooved Collagen Substrate Enabling the Modeling of an In Vivo Cell Arrangement. Bioengineering (Basel), 2021. 8(9).

34. Qi, Y., et al., RhoA/ROCK Pathway Activation is Regulated by AT1 Receptor and Participates in Smooth Muscle Migration and Dedifferentiation via Promoting Actin Cytoskeleton Polymerization. International Journal of Molecular Sciences, 2020. 21(15).

35. Lv, P., et al., SM22α inhibits lamellipodium formation and migration via Ras-Arp2/3 signaling in synthetic VSMCs. American Journal of Physiology-Cell Physiology, 2016. 311(5): p. C758–C767.

36. Sun, Z.Q., S.S. Guo, and R. Fässler, Integrin-mediated mechanotransduction. Journal of Cell Biology, 2016. 215(4): p. 445–456.

37. Lim, S.M., et al., RhoA-induced cytoskeletal tension controls adaptive cellular remodeling to mechanical signaling. Integrative Biology, 2012. 4(6): p. 615–627.

38. Lin, P.K. and G.E. Davis, Extracellular Matrix Remodeling in Vascular Disease: Defining Its Regulators and Pathological Influence. Arteriosclerosis Thrombosis and Vascular Biology, 2023. 43(9): p. 1599–1616.

39. Suna, G., et al., Extracellular Matrix Proteomics Reveals Interplay of Aggrecan and Aggrecanases in Vascular Remodeling of Stented Coronary Arteries. Circulation, 2018. 137(2): p. 166–183.

40. Bao, H., et al., Platelet-Derived Extracellular Vesicles Increase Col8a1 Secretion and Vascular Stiffness in Intimal Injury. Frontiers in Cell and Developmental Biology, 2021. 9.

41. Deffur, A., et al., ANIMA: Association network integration for multiscale analysis. Wellcome Open Res, 2018. 3: p. 27.

42. Ruiz, C., M. Zitnik, and J. Leskovec, Identification of disease treatment mechanisms through the multiscale interactome. Nature Communications, 2021. 12(1).

43. Xu, Z.H., et al., Development of Multiscale Transcriptional Regulatory Network in Esophageal Cancer Based on Integrated Analysis. Biomed Research International, 2020. 2020.

44. Kumar, R., et al., Differential gene expression and protein-protein interaction network profiling of sulfur mustard-exposed rabbit corneas employing RNA-seq data and bioinformatics tools. Experimental Eye Research, 2023. 235.

45. Avelar, R.A., et al., A multidimensional systems biology analysis of cellular senescence in aging and disease. Genome Biology, 2020. 21(1).

46. Xu, P. and B. Zhang, Multiscale network modeling reveals the gene regulatory landscape driving cancer prognosis in 32 cancer types. Genome Research, 2023. 33(10): p. 1806–1817.

47. Love, M.I., W. Huber, and S. Anders, Moderated estimation of fold change and dispersion for RNA-seq data with DESeq2. Genome Biology, 2014. 15(12).

48. Wickham, H., ggplot2: Elegant Graphics for Data Analysis. Ggplot2: Elegant Graphics for Data Analysis, 2009: p. 1–212.

49. Reimand, J., et al., g:Profiler -: a web-based toolset for functional profiling of gene lists from large-scale experiments. Nucleic Acids Research, 2007. 35: p. W193–W200.

50. Krämer, A., et al., Causal analysis approaches in Ingenuity Pathway Analysis. Bioinformatics, 2014. 30(4): p. 523–530.

51. Shannon, P., et al., Cytoscape: A software environment for integrated models of biomolecular interaction networks. Genome Research, 2003. 13(11): p. 2498–2504.

52. Bae, Y.H., et al., A FAK-Cas-Rac-lamellipodin signaling module transduces extracellular matrix stiffness into mechanosensitive cell cycling. Sci Signal, 2014. 7(330): p. ra57.

53. Krajnik, A., et al., Survivin regulates intracellular stiffness and extracellular matrix production in vascular smooth muscle cells. APL Bioeng, 2023. 7(4): p. 046104.

54. Di, X., et al., Cellular mechanotransduction in health and diseases: from molecular mechanism to therapeutic targets. Signal Transduct Target Ther, 2023. 8(1): p. 282.

55. Allen, A., D. Gau, and P. Roy, The role of profilin-1 in cardiovascular diseases. J Cell Sci, 2021. 134(9).

56. Regalado, E.S., et al., Aortic Disease Presentation and Outcome Associated With ACTA2 Mutations. Circ Cardiovasc Genet, 2015. 8(3): p. 457–64.

57. Negishi, K., et al., Author Correction: An Myh11 single lysine deletion causes aortic dissection by reducing aortic structural integrity and contractility. Sci Rep, 2024. 14(1): p. 7874.

58. Pucci, L., et al., A New Variant in the MYH11 Gene in a Familial Case of Thoracic Aortic Aneurysm. Ann Thorac Surg, 2020. 109(4): p. e279–e281.

59. Cruz-López, E.O., et al., Angiotensinogen Suppression: A New Tool to Treat Cardiovascular and Renal Disease. Hypertension, 2022. 79(10): p. 2115–2126.

60. Kobori, H., et al., Urinary angiotensinogen as a potential biomarker of severity of chronic kidney diseases. Journal of the American Society of Hypertension, 2008. 2(5): p. 349–354.

61. da Cunha, R.S., et al., Uremic toxins activate CREB/ATF1 in endothelial cells related to chronic kidney disease. Biochemical Pharmacology, 2022. 198.

62. Florczyk-Soluch, U., K. Polak, and J. Dulak, The multifaceted view of heart problem in Duchenne muscular dystrophy. Cell Mol Life Sci, 2021. 78(14): p. 5447–5468.

63. Lancioni, A., et al., Combined deficiency of alpha and epsilon sarcoglycan disrupts the cardiac dystrophin complex. Hum Mol Genet, 2011. 20(23): p. 4644–54.

64. Averill, M.M., C. Kerkhoff, and K.E. Bornfeldt, S100A8 and S100A9 in cardiovascular biology and disease. Arterioscler Thromb Vasc Biol, 2012. 32(2): p. 223–9.

65. Cotoi, O.S., et al., Plasma S100A8/A9 correlates with blood neutrophil counts, traditional risk factors, and cardiovascular disease in middle-aged healthy individuals. Arterioscler Thromb Vasc Biol, 2014. 34(1): p. 202–10.

66. Meng, Y., et al., Identification of Potential Key Genes Involved in the Carotid Atherosclerosis. Clin Interv Aging, 2021. 16: p. 1071–1084.

67. Yi, X., et al., Genetic variants of PTGS2, TXA2R and TXAS1 are associated with carotid plaque vulnerability, platelet activation and TXA2 levels in ischemic stroke patients. PLoS One, 2017. 12(7): p. e0180704.

